# Fast open modification spectral library searching through approximate nearest neighbor indexing

**DOI:** 10.1101/326173

**Authors:** Wout Bittremieux, Pieter Meysman, William Stafford Noble, Kris Laukens

## Abstract

Open modification searching (OMS) is a powerful search strategy that identifies peptides carrying any type of modification by allowing a modified spectrum to match against its unmodified variant by using a very wide precursor mass window. A drawback of this strategy, however, is that it leads to a large increase in search time. Although performing an open search can be done using existing spectral library search engines by simply setting a wide precursor mass window, none of these tools have been optimized for OMS, leading to excessive runtimes and suboptimal identification results. Here we present the ANN-SoLo tool for fast and accurate open spectral library searching. ANN-SoLo uses approximate nearest neighbor indexing to speed up OMS by selecting only a limited number of the most relevant library spectra to compare to an unknown query spectrum. This approach is combined with a cascade search strategy to maximize the number of identified unmodified and modified spectra while strictly controlling the false discovery rate, as well as a shifted dot product score to sensitively match modified spectra to their unmodified counterparts. ANN-SoLo achieves state-of-the-art performance in terms of speed and the number of identifications. On a previously published human cell line data set, ANN-SoLo confidently identifies more spectra than SpectraST or MSFragger and achieves a speedup of an order of magnitude compared to SpectraST.

ANN-SoLo is implemented in Python and C**++**. It is freely available under the Apache 2.0 license at https://github.com/bittremieux/ANN-SoLo.

## 1 Introduction

Although mass spectrometry (MS) is a very powerful technique to characterize proteins in complex biological samples, a significant portion of the thousands of spectra that are typically generated during a shotgun proteomics experiment cannot be confidently identified. In many cases a spectrum cannot be identified because it was generated by a peptide that contains one or more post-translational modifications (PTMs) [15]. If a particular modification has not been specified in the search settings, then spectra corresponding to peptides harboring this modification will be assigned an incorrect amino acid sequence. Because such false hits tend to receive higher scores than other false positive matches, these missed modifications have a detrimental effect on the identification performance [8].

On the one hand, protein modifications can be an artifact of the MS process because sometimes they are introduced during sample preparation [7]. A common example is alkylation by using iodoacetamide, which leads to the attachment of a carbamidomethyl group to cysteine residues, preventing the denatured proteins from reforming disulfide bridges. On the other hand, naturally occurring PTMs can be very interesting from a biological perspective as they often play important roles in many cellular processes [65].

Unfortunately, although MS techniques have become quite mature, comprehensive identification of modified proteins in complex samples remains challenging [1, 39]. In the traditional sequence database searching paradigm, all modifications of interest have to be explicitly specified in the search settings to correctly identify the spectra that include one or more of these modifications. This requirement leads to a significant search space increase because, for each peptide, both its unmodified version and all possible modified variants need to be considered, which results in an increased computational load and reduced sensitivity. Consequently, only a limited selection of the most prevalent modifications are commonly considered.

Alternatively, open modification searching (OMS) is a powerful strategy to identify modified spectra. Whereas traditionally only candidates that fall within a limited mass window around the query spectrum’s precursor mass are considered as a potential match, during OMS a very wide precursor mass window exceeding the delta mass induced by a PTM is used. This approach makes it possible to compare a modified query spectrum to its unmodified variant [2, 48]. As such, during an open search all possible protein modifications for which the mass difference falls within the precursor mass window, which is typically on the order of several hundreds of Dalton, are implicitly considered, including PTMs, amino acid substitutions, cleavage variants, etc. Afterwards, the presence and type of each modification can be derived from the difference between the observed precursor mass and the mass of the unmodified peptide.

Although OMS makes it possible to identify a wide range of spectra containing diverse modifications, the use of a very wide precursor mass window leads to a drastically increased search space. Consequently, compared to a standard search, the computational cost for an open search is orders of magnitude higher. A popular historical approach to keep OMS computationally feasible has been to use spectral libraries [3, 11, 34, 43, 67]. Because spectral libraries only contain previously observed peptides, the search space was often substantially restricted in comparison to sequence database searching strategies that consider all theoretically possible peptides [31]. On the other hand, with the increasing availability of high quality data sets in public data repositories nowadays [53], spectral libraries have grown substantially (figure 1). Indeed, for some well-studied organisms the spectral library size can rival the size of the sequence database. Consequently, scalable solutions for open spectral library searching are necessary to fully exploit the current wealth of information contained in large spectral libraries.

**Figure 1:**
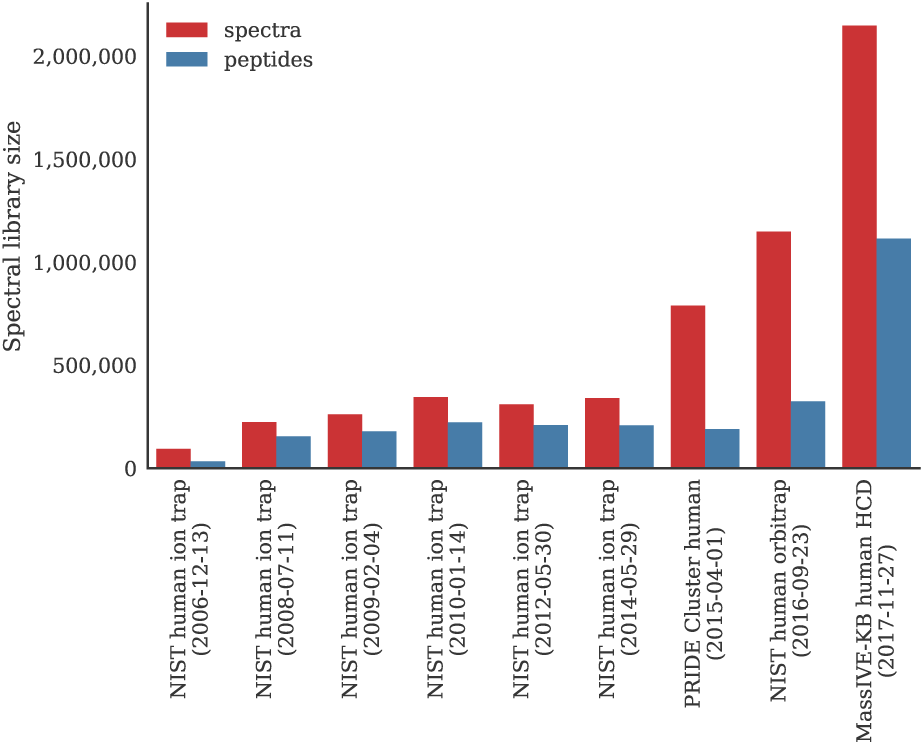
The size of spectral libraries has increased as more high-quality data sets have become available in public data repositories. Whereas traditionally spectral libraries were explicitly curated and compiled, e.g. by the National Institute of Standards and Technology (NIST), a recent alternative has been to automatically generate large spectral libraries on a repository-wide scale, e.g. based on all data sets in the PRoteomics IDEntifications (PRIDE) and MassIVE databases [32, 33].

Here we present the **A**pproximate **N**earest **N**eighbor **S**pectral **L**ibrary (ANN-SoLo) search tool, which has been optimized for fast and accurate open modification spectral library searching. Using a cascade search strategy [38] ANN-SoLo first identifies any unmodified peptides, followed by an open search to identify the modified peptides. During this open search ANN-SoLo uses an approximate nearest neighbor (ANN) index to efficiently find a limited set of the most similar library spectra for each query spectrum. During the open search the shifted dot product is used to accurately match modified query spectra to their unmodified library counterpart by taking into account peak shifts caused by a modification [11, 34].

Although some approaches have previously been proposed to speed up spectral library searching, including parallelizing spectral matching using graphics processing units (GPUs) [6] and candidate filtering based on a shared peak count [64], these approaches did not deal with the specific challenges posed by OMS. In contrast, the Liberator and MzMod software tools are based on the Apache Spark framework for distributed data processing to massively parallelize open spectral library searching [34]. Although these big data tools make it possible to process large amounts of spectral data, they require a specialized cluster or cloud infrastructure.

Additionally, several software tools have recently been developed to speed up OMS when using a sequence database instead of a spectral library. MSFragger uses an index of theoretical fragments to quickly compute the number of shared fragment ions between a query spectrum and theoretical spectra [40]. SpecOMS uses an FPtree-like data structure [9], called SpecTrees, to encode the number of shared masses between all spectra [17]. Sequence tags are a popular approach to restrict the search space as well. PIPI uses sequence tags of length 3 to perform a fuzzy tag-based filtering [68]. TagGraph uses an FM-index to filter candidates based on substrings of *de novo* derived sequences [20]. Finally, Open-pFind combines tag-based filtering to speed up open searches with a subsequent standard search including highly abundant variable modifications in a two-pass strategy [14].

Generally MS/MS spectrum identification can be considered a nearest neighbor task: for a given query spectrum the most similar database spectrum, i.e. its nearest neighbor, be it a real spectrum during spectral library searching or a theoretical spectrum during sequence database searching, has to be retrieved. Consequently these nearest neighbor queries can be sped up by making use of index structures. For example, a multiple vantage point tree [54] and locality-sensitive hashing (LSH) [22] have previously been proposed to speed up sequence database searching.

These various tools that have recently been developed for efficient OMS on the one hand and the application of multidimensional indexing techniques to speed up spectral identification on the other hand have so far exclusively focused on sequence database searching. Here, we show how ANN indexing can be used to speed up open spectral library searching. Our ANN-SoLo tool is able to efficiently search very large spectral libraries and sensitively identify spectra containing any modification, outperforming other spectral library search engines in both speed and the number of identified spectra.

ANN-SoLo is implemented in Python and C**++**. It is freely available as open source under the permissive Apache 2.0 license at https://github.com/bittremieux/ANN-SoLo.

## 2 Methods

### 2.1 Cascade spectral library searching

Spectral library searching works by comparing experimental, unknown query spectra to previously observed, known spectra in the spectral library. To identify a query spectrum, its best matching library spectrum is found and is assigned the corresponding peptide sequence [31, 57]. Finding the closest matching library spectrum for a given query spectrum can be divided into two steps: (i) a candidate selection step during which a subset of spectra in the spectral library are selected as candidate matches, and (ii) a candidate ranking step during which, for each candidate spectrum, a spectrum–spectrum match (SSM) score is calculated to quantify the similarity between the two spectra. Subsequently the candidate match with the highest score is used as the identification for the query spectrum.

To perform an open search ANN-SoLo employs a cascade search strategy consisting of two levels, which allows it to maximize the number of identified spectra while strictly controlling the false discovery rate (FDR) [38]. In the first level of the cascade search a small precursor mass window is used to identify any unmodified spectra. The resulting SSMs are filtered on FDR, and the confident SSMs below the FDR threshold are retained. Next, the SSMs exceeding the FDR threshold are passed on to the second level of the cascade search, in which a wide precursor mass window is used to identify modified spectra. The resulting SSMs are filtered on FDR as well and combined with the accepted SSMs from the first level to form the final set of spectrum identifications.

Below, we describe how ANN-SoLo addresses the candidate selection and candidate ranking steps during both levels of the cascade open search, and describe how it achieves both speed and accuracy.

#### 2.1.1 Spectrum preprocessing

Prior to spectral library searching both the query spectra and library spectra are similarly preprocessed to represent the spectra in a uniform way and discard low-quality spectra [55]. Peaks corresponding to the precursor ion and noise peaks with an intensity below 1 % of the intensity of the most intense peak are removed and, if applicable, the spectrum is further restricted to its 50 most intense peaks [42]. After peak removal, any spectrum that contains fewer than 10 peaks remaining or with a mass range less than 250 Da is discarded. Finally, peak intensities are rank transformed to de-emphasize overly dominant peaks [43].

#### 2.1.2 Candidate selection

Typically, the candidate selection step consists of a precursor mass filter, i.e. only the library spectra whose precursor mass falls within a narrow window around the query spectrum’s precursor mass are considered as candidates. Especially for modern high-resolution instruments, which can report masses with a (sub-)ppm accuracy, the number of considered candidate spectra can be very small. However, when a wider precursor mass window is used, as in the case of open searches, the number of candidates that are selected can increase by several orders of magnitude. In this case the precursor mass window will not be an effective filter, and the search time will increase accordingly.

During the first level of its cascade search ANN-SoLo uses a small precursor mass window, and the search proceeds in the standard fashion. In contrast, during the second level of its cascade search ANN-SoLo uses an ANN index consisting of an ensemble of random projection trees [5] to efficiently filter the library spectra based on their similarity to the query spectra.

To construct the ANN index each library spectrum is vectorized to represent it as a point in a multidimensional space. A spectrum is converted into a sparse vector by dividing it into mass bins of 1 Da and assigning its peak intensities to their corresponding mass bins, after which the vector is normalized to have unit length. In case multiple peaks in the mass spectrum are assigned to the same mass bin their intensities are summed. For low mass accuracy spectra, this procedure will occasionally incorrectly assign peaks to neighboring bins. However, the binning procedure is only applied during candidate selection, which is robust to mismatched bin assignments because a sufficiently high number of candidates are retrieved from the ANN index. Next, the vectors for all library spectra are used to build a binary index tree (figure 2). This is done by recursively partitioning the data space into two subspaces using random split hyperplanes. Concretely, two points are randomly sampled to construct a split hyperplane equidistant from both points. This hyperplane divides the data points into two subspaces based on their position relative to it (figure 2a). Next, for each of these two subspaces the same procedure can be repeated: by randomly drawing a new split hyperplane into the data subspace it can be further partitioned into two smaller subspaces. This process is recursively repeated to construct a binary index tree (figures 2b and 2c).

**Figure 2:**
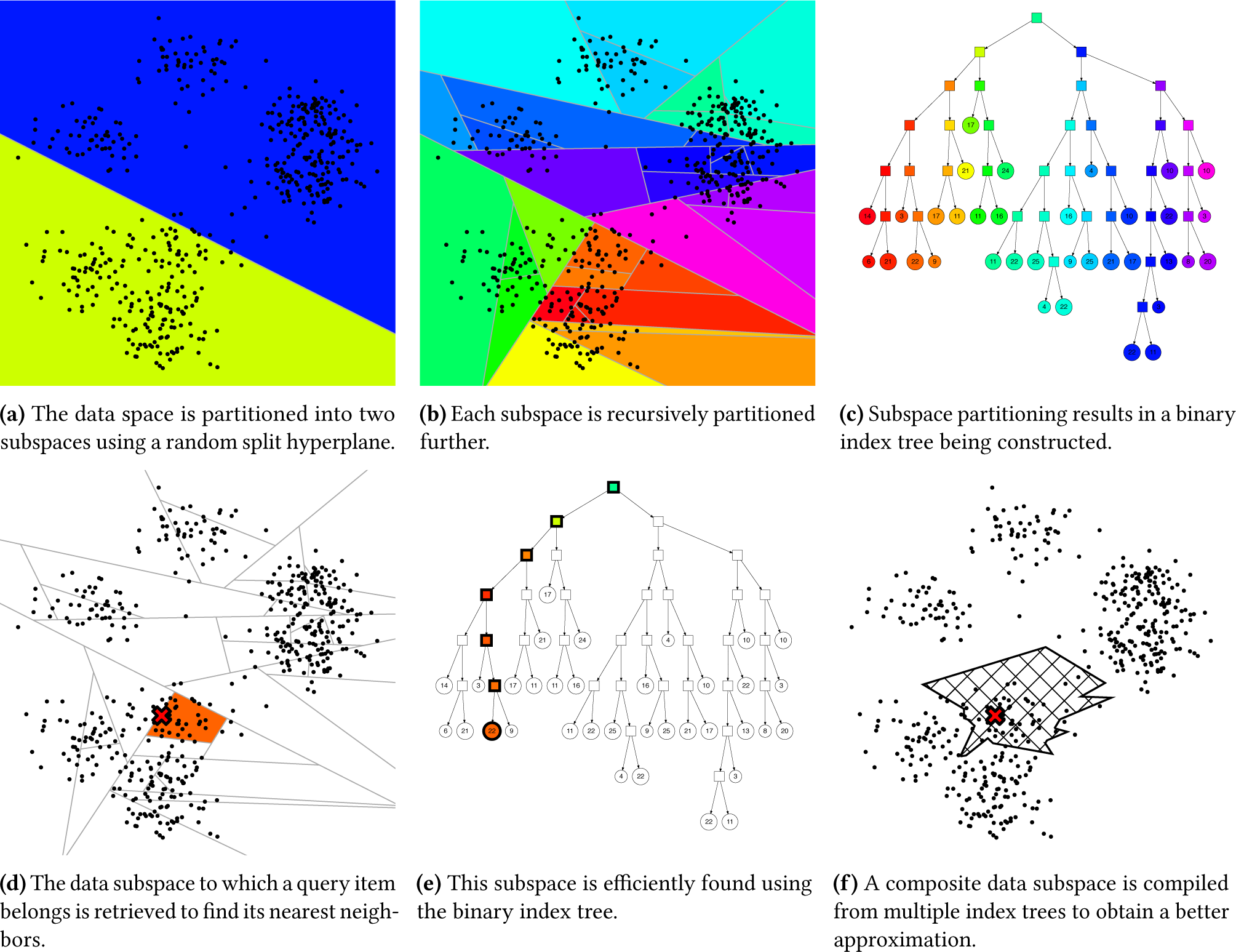
Approximate nearest neighbor indexing and searching using an ensemble of random split hyperplane index trees.

This binary index tree can be used to efficiently find the nearest neighbor for a given query point. Instead of having to compare the query point to all data points the index tree can be traversed to find the data subspace to which the query point would belong and which will likely contain its nearest neighbor (figures 2d and 2e). In this fashion a nearest neighbor query can be performed in logarithmic time in terms of the number of data points, whereas it would require linear time in terms of the number of data points to compare the query point against all data points in a brute-force fashion.

Unfortunately, the nearest neighbor for the query point might not be located in the data subspace that has been selected, but instead it might be present in an adjacent data subspace. In this case it is not possible to directly identify the actual nearest neighbor and only an approximate result will be achieved. To reduce the risk of missing the actual nearest neighbor multiple complementary index trees are used. Because random split hyperplanes are used to divide the data space while constructing the initial tree, an alternative index tree can be created by making use of different split hyperplanes. The new random split hyperplanes cause the data space to be subdivided differently, leading to a different binary index tree. As a result, the risk of missing an actual nearest neighbor is minimized by using both index trees simultaneously to answer a query. During querying the data subspaces to which a query point belongs are identified for each tree individually, after which the data points in both of these subspaces are combined to find the actual nearest neighbor. Finally, additional index trees can be constructed in a similar fashion to form an ensemble of index trees, with each tree providing a complementary view on the data. Through the combination of this ensemble of index trees the risk of missing the actual nearest neighbor is further decreased (figure 2f). As such the number of trees in the ensemble is a hyperparameter that can be used to configure a trade-off between accuracy and speed, as using more trees reduces the risk of missing the actual nearest neighbor at the expense of some increased computational requirements.

#### 2.1.3 Candidate ranking

During the candidate ranking step the similarities between the query spectrum and all library spectra that have been selected in the previous step are evaluated to determine the highest-scoring SSM. Even though the ANN index already retrieves the most likely candidate spectra based on their similarity, a subsequent ranking step remains necessary. This is because the vectors employed for ANN indexing only represent the spectra at a coarse, 1 Da bin granularity. In contrast, for highresolution mass spectra a more accurate score can be computed using a low fragment mass tolerance to obtain the optimal match. Additionally, during the open search the shifted dot product is used as scoring method to take PTMs into account and accurately match modified spectra to their unmodified variant, as described next.

The dot product is a well-established scoring method to rank SSMs. An important advantage of the dot product is that, despite its simplicity, it is able to accurately capture the similarity between two mass spectra [60]. Additionally, it can be computed very efficiently as it has a time complexity of *O*(*n*), where *n* is the total number of peaks in the two spectra being compared. Based on these advantageous properties the dot product has been used by several spectral library search engines [25, 42]. Similarly, ANN-SoLo uses the dot product to identify unmodified spectra during the first level of its cascade search.

However, because the dot product only considers directly matching peaks with identical masses (while taking the fragment mass tolerance into account) it is less suitable to identify modified spectra. Instead, during the second level of its cascade search ANN-SoLo uses a variation on the dot product, called the shifted dot product [11, 34]. This shifted dot product also considers peaks that are shifted according to the precursor mass difference between the two spectra that are being matched to accurately identify modified spectra (figure 3). We here briefly describe an algorithm to compute the shifted dot product in *O*(*n* log *n*) time complexity [34].

**Figure 3:**
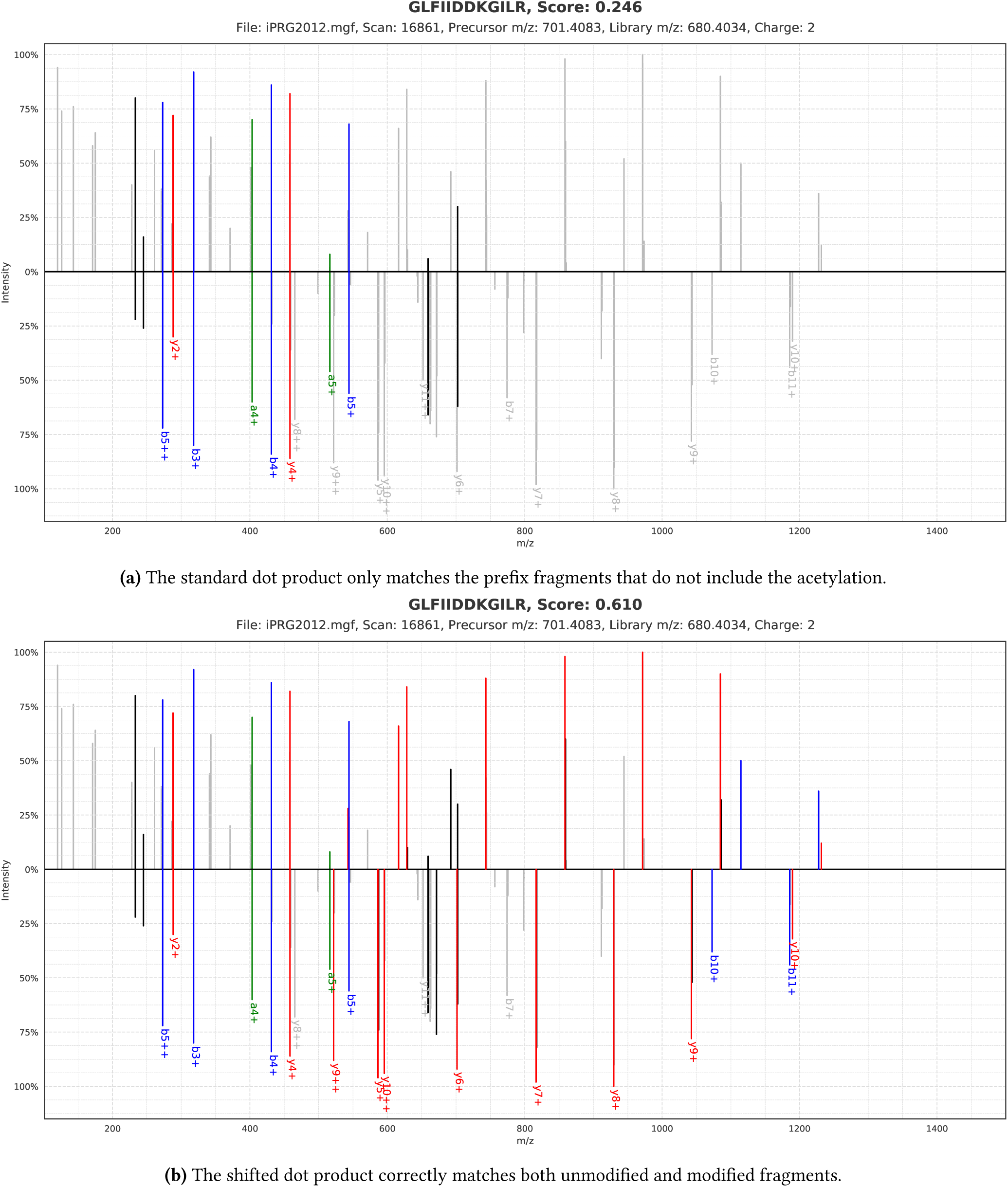
The shifted dot product enables more accurate matching between an unmodified library spectrum (bottom) and a modified query spectrum (top) than the standard dot product. As can be derived from the precursor mass difference the peptide GLFIIDDKGILR has undergone acetylation (mass 42.010 565 Da) on the lysine at position 8. The standard dot product only takes directly matching peaks into account, while the shifted dot product can consider shifted peaks according to the precursor mass difference and charge, correctly assigning a high score to this match.

First, the precursor mass difference between the two spectra is calculated and normalized according to the precursor charge. Next, all potential peak pairs, with and without a mass shift (taking into account different charges), can be determined in a linear pass through both spectra. Each peak pair is scored by multiplying the intensities of both peaks, as in the standard dot product. Unshifted peak matches and shifted peak matches that include an annotated peak are scored fully while shifted peak matches without an annotation are slightly penalized to minimize the influence of potential spurious matches. Next, to calculate the final shifted dot product score the peak matches are sorted on their intensity product, after which they are summed in a greedy fashion while avoiding to match a single peak in either of the two spectra more than once. Because peak matches are selected individually, this heuristic approach to calculate the shifted dot product does not explicitly try to ensure that a consecutive range of fragment peaks along the peptide sequence are consistently shifted. This approach allows us to efficiently select all peak matches using only a single simultaneous pass through both spectra, whereas a full spectrum alignment would require *O*(*n*^2^) time complexity in the number of peaks.

### 2.2 FDR calculation

False discovery rates are calculated in two phases after both levels of the cascade search using the target–decoy strategy [23]. Specifically, FDRs are estimated based on a concatenated spectral library containing both target and decoy spectra (see section 2.3) using the number of decoys divided by the number of target SSMs.

During the first level of the cascade search the SSMs are directly filtered on the FDR. In contrast, because the score distributions of peptides with different modifications can exhibit distinct properties, during the second level of the cascade search the SSMs are filtered using the subgroup FDR strategy [26, 27]. To combine SSMs with identical modifications they are grouped based on their charge-normalized precursor mass difference between the query spectra and library spectra. The SSMs are split into subgroups by iteratively selecting the SSM with the highest match score alongside all other remaining SSMs whose precursor mass difference falls within a 0.1 Da range. Because FDR estimates become progressively less reliable if only a limited number of observations are considered FDR filtering is only done for subgroups that contain at least 20 SSMs. Subgroups that contain fewer SSMs are combined into a residual group instead whose FDR is jointly calculated in the end.

### 2.3 Data sets

The first data set we used was generated in the context of the 2012 study by the Proteome Informatics Research Group of the Association of Biomolecular Resource Facilities. The goal of this study was to assess the community’s ability to analyze modified peptides [12]. Towards this end, various participating researchers were asked to identify an unknown data set, after which their proficiency in handling modified peptides was evaluated. The provided data set consisted of a mixture of synthetic peptides with biologically occurring modifications combined with a yeast whole cell lysate as background, and the spectra were measured using a TripleTOF instrument. For full details on the sample preparation and acquisition see the original publication by Chalkley et al. [12]. This high quality data set has been recommended as a reference data set for the evaluation of identification algorithms [28]. All data was downloaded from the MassIVE data repository (accession MSV000078492).

To search the iPRG2012 data set the human HCD spectral library compiled by NIST (version 2016/09/12) and a TripleTOF yeast spectral library from Selevsek et al. [56] were used. First, matches to decoy proteins were removed from the yeast spectral library, after which both spectral libraries were concatenated using SpectraST [42] while removing duplicates by retaining only the best replicate spectrum for each individual peptide ion. Next, decoy spectra were added in a 1:1 ratio using the shuffle-and-reposition method [41], resulting in a single large spectral library file containing 1 188 168 spectra.

The second data set consists of spectra measured from the HEK293 human cell line [15]. As per Chick et al. [15], the HEK293 cells were first lysed, trypsinized, and separated into 24 fractions, after which high-resolution and high-mass accuracy MS/MS spectra were obtained on an LTQ Orbitrap Elite mass spectrometer. For full details on the sample preparation and acquisition see the original publication by Chick et al. [15]. Raw files were downloaded from PRIDE [63] (project PXD001468) and converted to MGF files using msconvert [13].

To search the HEK293 data set the MassIVE-KB peptide spectral library (version 2017/11/27) was used. This is a repository-wide human higher-energy collisional dissociation spectral library derived from over 30 TB of human MS/MS proteomics data. The original spectral library contained 2 148 752 MS/MS spectra, from which duplicates were removed using SpectraST [42] by retaining only the best replicate spectrum for each individual peptide ion, resulting in a spectral library containing 1 504 951 spectra. Next, decoy spectra were added in a 1:1 ratio using the shuffle-and-reposition method [41], resulting in a final spectral library containing 3 009 902 spectra. To the best of our knowledge this is the largest spectral library reported in the literature to date.

All MS/MS data, spectral libraries, and identification results have been deposited to the ProteomeXchange Consortium [19] via the PRIDE partner repository [63] with the data set identifier PXD009861.

### 2.4 Search settings

#### 2.4.1 ANN-SoLo

The spectrum preprocessing settings are as described in section 2.1.1. For the iPRG2012 data set a precursor mass window of 20 ppm was used for the standard search and 20 ppm followed by 300 Da for the cascade open searches. Additionally, a fragment mass tolerance of 0.02 Da was used in all cases. For the HEK293 data set a precursor mass window of 5 ppm was used for the standard searches and a precursor mass window of 5 ppm followed by 500 Da was used for the cascade open searches. For all HEK293 searches a fragment mass tolerance of 0.02 Da was used.

Two major hyperparameters influence the performance of the ANN index: the number of trees in the index and the number of nodes to inspect per query. We discuss the effect of these hyperparameters in detail in section 3.2. For the iPRG2012 data set we used ANN indices consisting of 100 to 1000 trees, while the number of nodes inspected during querying was varied between 20 000 and 400 000. To analyze the HEK293 data set an ANN index consisting of 1000 trees was used and 200 000 nodes were inspected during searching.

#### 2.4.2 SpectraST

We compared the performance of ANN-SoLo against the popular spectral library search engine SpectraST [42].

We used SpectraST version 5.0 as part of the TransProteomic Pipeline version 5.1.0 [18]. We tried to specify the SpectraST search settings for processing the HEK293 data set as closely as possible to the ANN-SoLo settings to ensure a fair comparison. Spectra were preprocessed to have a minimum mass range of 250 Da and only the 50 most intense peaks were retained. Peak-to-peak matching and rank-based scoring was used to evaluate SSMs. Library caching was enabled, which is an essential requirement in order to be able to complete the open searches. Similar to the ANN-SoLo settings a precursor mass window of 500 Da was used for the open searches. In contrast, for the standard searches a precursor mass window of 0.02 Da was used as SpectraST does not support specification of tolerances in ppm units. A fragment mass tolerance of 0.02 Da was used in all cases. FDRs were estimated in a post-processing step using the subgroup FDR strategy described in section 2.2.

#### 2.4.3 MSFragger

We also compared the performance of ANN-SoLo against the recent state-of-the-art OMS sequence database search engine MSFragger [40].

We used MSFragger version 2017/11/06. Because MSFragger is a sequence database search engine, whereas ANN-SoLo and SpectraST are both spectral library search engines, settings for MSFragger necessarily slightly deviate. To search the HEK293 data set we used the neXtProt database of human protein sequences [29] (version 2018/01/17) complemented with common contaminants from the cRAP database (version 2012/01/01). Next, a concatenated target–decoy database was generated by appending an equal number of shuffled decoy sequences using Crux [45]. MSFragger was configured to consider tryptic peptides with up to 2 missed cleavages, and cysteine carbamidomethylation was specified as a static modification. A precursor mass window of 5 ppm was used for the standard searches, and a precursor mass window of 500 Da was used for the open searches. In both cases a fragment mass tolerance of 0.02 Da was used. FDRs were estimated in a post-processing step using the subgroup FDR strategy described in section 2.2.

### 2.5 Code availability

The ANN-SoLo software is written in Python, making use of various open-source libraries such as NumPy [62], SciPy, and pandas [46] for scientific computing and Matplotlib [35], Seaborn [66], and Jupyter notebooks [52] for visualization purposes. Pyteomics [30] is used to support some mass spectrometry-specific functionality, such as reading input files and for FDR calculation.

ANN indexing during the candidate selection step is based on the open-source Approximate Nearest Neighbors Oh Yeah (Annoy) library [59], which was originally developed at Spotify to support large-scale music recommendations. Dot product and shifted dot product calculation is implemented as an external C**++** module to optimize the candidate ranking step for speed.

All code is released as open source under the permissive Apache 2.0 license and is available at https://github.com/bittremieux/ANN-SoLo. This web resource also includes detailed instructions on how to install and run ANN-SoLo, along with code notebooks to reproduce all analyses discussed next.

## 3 Results

### 3.1 Cascade open search maximally identifies unmodified and modified peptides

The advantage of an open search compared to a standard search is that modified peptides can be identified without having to specify the expected modifications. However, using a wide precursor mass window might lead to a loss in identifications of spectra that would have previously been identified during a standard search. Whereas a specific query spectrum might be identified differently with a slightly higher score when using a wide precursor mass window than when using a small precursor mass window, as the search space is significantly larger in the former case these scores cannot be directly compared. Instead, through its cascade search strategy ANN-SoLo maximally identifies both unmodified and modified peptides. Based on the iPRG2012 data set ANN-SoLo achieves a 25 % increase in identifications when performing an open search compared to a standard search (table 1), with the additional identifications corresponding to modified peptides.

Moreover, a further 21 % increase in identifications is achieved by using the shifted dot product instead of the standard dot product to score the spectral similarity of modified spectra (table 1). Because the shifted dot product explicitly accounts for modification-induced shifts in the fragment ion peaks it more accurately identifies modified spectra. As a result, by employing a cascade open search strategy and using an optimized scoring function modified spectra can be accurately matched to the spectral library, resulting in a total increase in identifications of 45 % when comparing a standard search against an open search on the iPRG2012 data set.

**Table 1:** The number of accepted SSMs at a 1 % FDR threshold for various searches of the iPRG2012 data set. The open search identifies all unmodified spectra previously identified in the standard search as well as additional modified spectra. A further increase in identifications is achieved by using the shifted dot product to score modified SSMs.

| Search mode | Similarity measure | # SSMs |
| --- | --- | --- |
| Standard search | dot product | 4141 |
| Open search | dot product | 5160 |
| Open search | shifted dot product | 6019 |

These identifications can be compared to the consensus identifications compiled from the submissions to the iPRG2012 study [12], which can be considered as the “ground truth” (figure 4). ANN-SoLo agrees with the iPRG2012 identifications for 70 % of its spectrum assignments (4193 SSMs). Of the other ANN-SoLo identifications (1826 SSMs) that do not match the iPRG2012 consensus results 735 confidently identified SSMs conflict with the iPRG2012 consensus results and 1091 SSMs are uniquely contributed by ANN-SoLo. Although the conflicting SSMs are likely misidentified by ANN-SoLo, a third of those SSMs (275 SSMs) show a very high sequence similarity between the peptide assigned by ANN-SoLo and the iPRG2012 consensus identification (edit distance at most 3 amino acids; supplementary figure S1). Therefore, although these identifications do not directly match the iPRG2012 consensus results they can still be considered to be correct in the context of an open search as they correspond to peptide sequences that differ only by prefix or postfix amino acids, caused by missed cleavages, or single amino acid substitutions. Indeed, 450 peptides from the consensus identifications corresponding to 648 conflicting SSMs do not appear in the spectral library, preventing these spectra from being correctly identified by ANN-SoLo. Instead, ANN-SoLo provides a partially correct peptide assignment, which demonstrates the power of open modification searching to circumvent the inherently limited coverage of spectral libraries.

**Figure 4:**
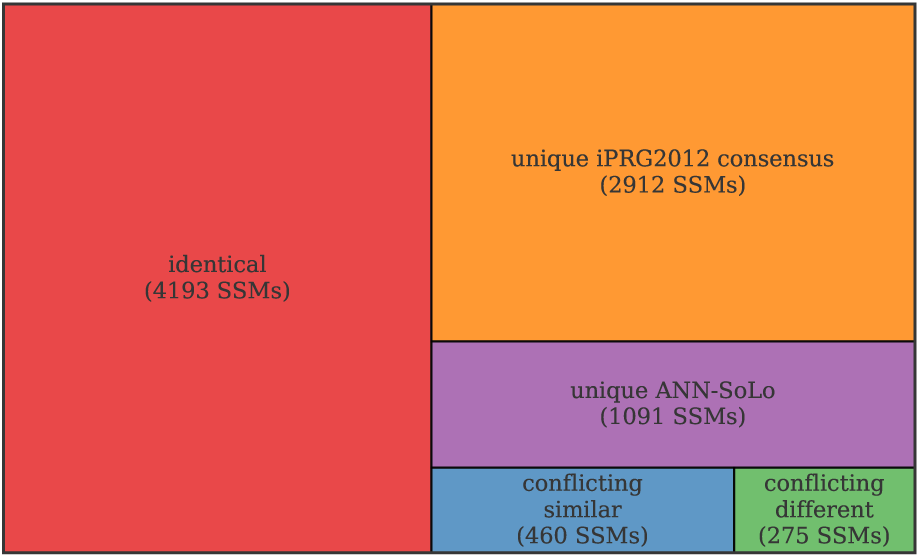
Agreement in SSMs between ANN-SoLo and the iPRG2012 consensus results.

Additionally, the advantage of the cascade search strategy over a direct open search can be demonstrated based on the iPRG2012 consensus ground truth. During the first stage of the cascade search only unmodified peptides are identified. Because we expect those peptides to occur more often than their modified variants, additional statistical power is gained by using a small precursor mass window in the first search iteration [49]. In contrast, when a wide precursor mass window is used directly some of the SSMs that are otherwise accepted during the first cascade stage might obtain a higher-scoring match against another library spectrum that was originally not considered, indicating potential false positive identifications (supplementary figure S2). ANN-SoLo easily allows the user to perform a direct open search as well by setting a wide precursor mass window for the first cascade stage and disabling the second cascade stage. On the iPRG2012 data set 690 SSMs receive a conflicting identification between the cascade search and the direct open search. As before, many of these conflicting identifications correspond to peptides that only differ by a small number of amino acids (supplementary figure S3). We can verify whether the cascade search or the direct open search provided the correct identification by comparing their conflicting SSMs to the iPRG2012 consensus ground truth. For 121 SSMs of the 690 conflicting SSMs no identification was available in the consensus results, while neither of the two strategies identified the completely correct peptide for another 88 spectra. For the remaining 481 spectra the cascade strategy identified the fully correct peptide sequences 89 % of the time (429 SSMs) compared to 11 % (52 SSMs) for the direct open search. For the nonoverlapping identifications between the cascade search and the direct open search, the cascade search correctly identifies more SSMs as well. While only 10 out of the 150 unique SSMs from the direct open search match the consensus results, for the cascade search 604 out of 777 unique SSMs match the consensus results. These results clearly demonstrate the advantage of the cascade search strategy, which allows ANN-SoLo to maximally identify both unmodified and modified peptides.

### 3.2 Approximate nearest neighbor indexing speeds up open search

By using an ANN index only a limited number of the most relevant library spectra are retrieved during the candidate selection step, which speeds up the subsequent candidate ranking step because far fewer SSM scores have to be computed. Two major hyperparameters influence the performance of the ANN index: the number of trees in the index ensemble, which controls the index construction during an initialization phase, and the number of tree nodes to inspect when evaluating a query, which controls the performance during searching.

Using more trees has a positive effect on the accuracy of the ANN index by providing multiple complementary views on the data subspaces, as detailed previously. However, this improvement comes at the expense of an increased memory consumption as the size of the ANN index scales linearly with the number of index trees. The number of tree nodes to inspect when evaluating a query can be used to configure the accuracy of the ANN index. Instead of only inspecting the subspace that exactly contains the query point the neighboring subspaces that are closest to the query point across all index trees can be inspected as well. This approach helps to avoid missing the nearest neighbors for the query point. Because the number of nodes to inspect is specified across all trees in the ANN index, this hyperparameter is to some extent related to the number of trees used.

These two hyperparameters constitute a trade-off between speed and accuracy, both when constructing the ANN index and at runtime. During querying, in general, using more trees and inspecting more nodes will lead to a more faithful approximation of the set of nearest neighbors retrieved from the ANN index, at the expense of an increase in computational requirements. Based on the iPRG2012 data set there is a clear difference in runtime between a standard search and the traditional brute-force approach of performing an open search on the one hand, and the advantage of using an ANN index for the open search on the other hand (figure 5 and supplementary table S1). As shown previously, the open search allows us to identify a significantly higher number of spectra, with the newly identified spectra corresponding to modified peptides. Unfortunately, this comes at the expense of a large increase in runtime, rendering OMS infeasible in practice. In contrast, by making use of an ANN index ANN-SoLo significantly decreases the time required to perform an open search, making OMS a viable strategy.

The ANN-SoLo speedup results from a reduction in the number of candidates that have to be evaluated during the candidate ranking step (supplementary figure S4). A standard search only takes a very short amount of time, with I/O costs for reading the experimental and library spectra forming the major bottleneck. In contrast, the massive increase in runtime of a brute-force open search is caused by the fact that when a very wide precursor mass window is used each query spectrum must be compared against a very large number of library spectra. Using an ANN index instead puts the focus on the candidate selection step to only retrieve the most relevant candidates. As a result, the relative proportion of work during the candidate ranking step significantly decreases. The ANN index also makes it possible to use complex scoring functions without incurring an overly excessive slowdown. Although the shifted dot product is computationally more expensive than the standard dot product, negative effects on the total runtime are limited due to the optimized candidate selection step.

**Figure 5:**
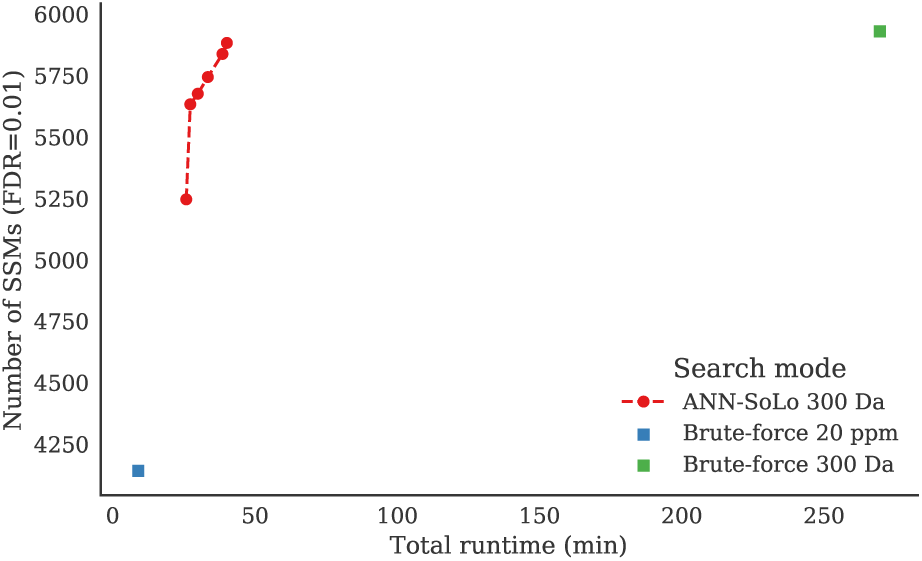
Runtime versus number of identifications for various searches of the iPRG2012 data set. Timing results in this figure and in table 2 were obtained on a single-core Intel Xeon E5-2680 v2 processor. An open search identifies a significantly higher number of spectra than a standard search at the expense of a large increase in runtime. The ANN-SoLo results show that the ANN index significantly reduces the time required for open searches. The multiple ANN-SoLo results correspond to different configurations of the ANN index hyperparameters, with the settings that lie on the Pareto frontier shown. Even when maximizing accuracy to achieve the same number of identifications as the brute-force approach ANN-SoLo considerably speeds up the search. Specific values for the ANN hyperparameters and the corresponding identification performance are available in supplementary table S1.

The previous timing results do not include the time required to construct the ANN index during an initial preprocessing phase. Especially when a large number of index trees are used in combination with a large spectral library, building the ANN index may take a non-negligible amount of time (supplementary table S1). However, this step only needs to happen once, after which the ANN index can be reused for many subsequent searches. Additionally, the space requirement of the ANN index is proportional to the size of the spectral library for which it is constructed and to the number of trees in the ANN ensemble. Although the ANN index will typically be larger than other commonly used indexing methods to assist MS/MS spectrum identification, such as a peptide index [21, 50] or a fragment ion index [40], because the ANN index contains a low-resolution vector representation for each spectrum in the spectral library, in general the memory requirements remain feasible for modern workstations. For efficient memory management ANN-SoLo splits the ANN index into individual files for different charge states and only processes a single file at a time, while using memory mapping to efficiently read the index files.

**Table 2:** Runtime and identification rates for ANN-SoLo, SpectraST, and MSFragger on the HEK293 data set. The runtime is reported in minutes and represents the average runtime over all 24 raw files. The identification rate is reported in terms of the number of accepted SSMs at 1 % FDR and the number of corresponding unique peptides and is reported for the entire 24-run data set.

| Search engine | Time (min) | # SSMs | # Peptides |
| --- | --- | --- | --- |
| *Standard search* |  |  |  |
| MSFragger | 0.7 | 344 998 | 104 672 |
| SpectraST | 5.2 | 369 079 | 102 077 |
| ANN-SoLo | 24.0 | 352 938 | 105 870 |
| *Open search* |  |  |  |
| MSFragger | 34.7 | 526 027 | 126 364 |
| SpectraST | 1276.7 | 473 729 | 112 375 |
| ANN-SoLo | 108.5 | 647 469 | 153 605 |

### 3.3 ANN-SoLo achieves state-of-the-art performance in terms of speed and number of identifications

We compared ANN-SoLo to SpectraST, a commonly used spectral library search engine, and MSFragger, a recent sequence database search engine optimized for OMS, in terms of speed and the number of identifications on the HEK293 data set (table 2). When comparing both spectral library search engines, for a standard search SpectraST is clearly faster. This can be attributed to simple implementation differences, of which a notable factor is that ANN-SoLo is mainly implemented in the Python programming language while SpectraST is implemented in C**++**. As Python is an interpreted programming language it can be up to a hundred times slower than a compiled programming language such as C**++**, as benchmarks have shown for several general tasks. In contrast, for the open search ANN-SoLo is an order of magnitude faster than SpectraST, reducing the search time from almost a day on average to under two hours, despite the inherent programming language disadvantage. This result clearly shows the massive advantage of ANN indexing to speed up OMS. In contrast, MSFragger is clearly faster than ANN-SoLo for both the standard search and the open search, showing its high performance in terms of search speed in general and the efficiency of its fragment ion indexing to speed up OMS.

In terms of peptide identifications, all three search engines show a high degree of agreement as well as complementarity (figure 6). For the standard search, each search engine provides a unique set of identifications, confirming the complementarity of different search strategies [58]. However, because ANN-SoLo and SpectraST use the same spectral library and the dot product to score SSMs, their peptide identifications from the standard search overlap to a significant extent. Meanwhile MSFragger benefits from a more comprehensive search space: because it uses a sequence database containing the entire human proteome instead of the spectral library, MSFragger is able to provide additional peptide identifications. The difference in identified peptides is more pronounced for the open search. Again, we observe considerable agreement among all three search engines. Additionally, ANN-SoLo and SpectraST show their agreement by sharing a significant number of peptide identifications not found by MSFragger. Furthermore, both ANN-SoLo and MSFragger provide a similar number of unique peptide identifications. In contrast, because SpectraST is not optimized for identifying modified peptides its contribution in terms of unique peptide identifications is minimal. These results indicate the beneficial performance of ANN-SoLo against the current state of the art in spectral library searching as well as open modification searching.

**Figure 6:**
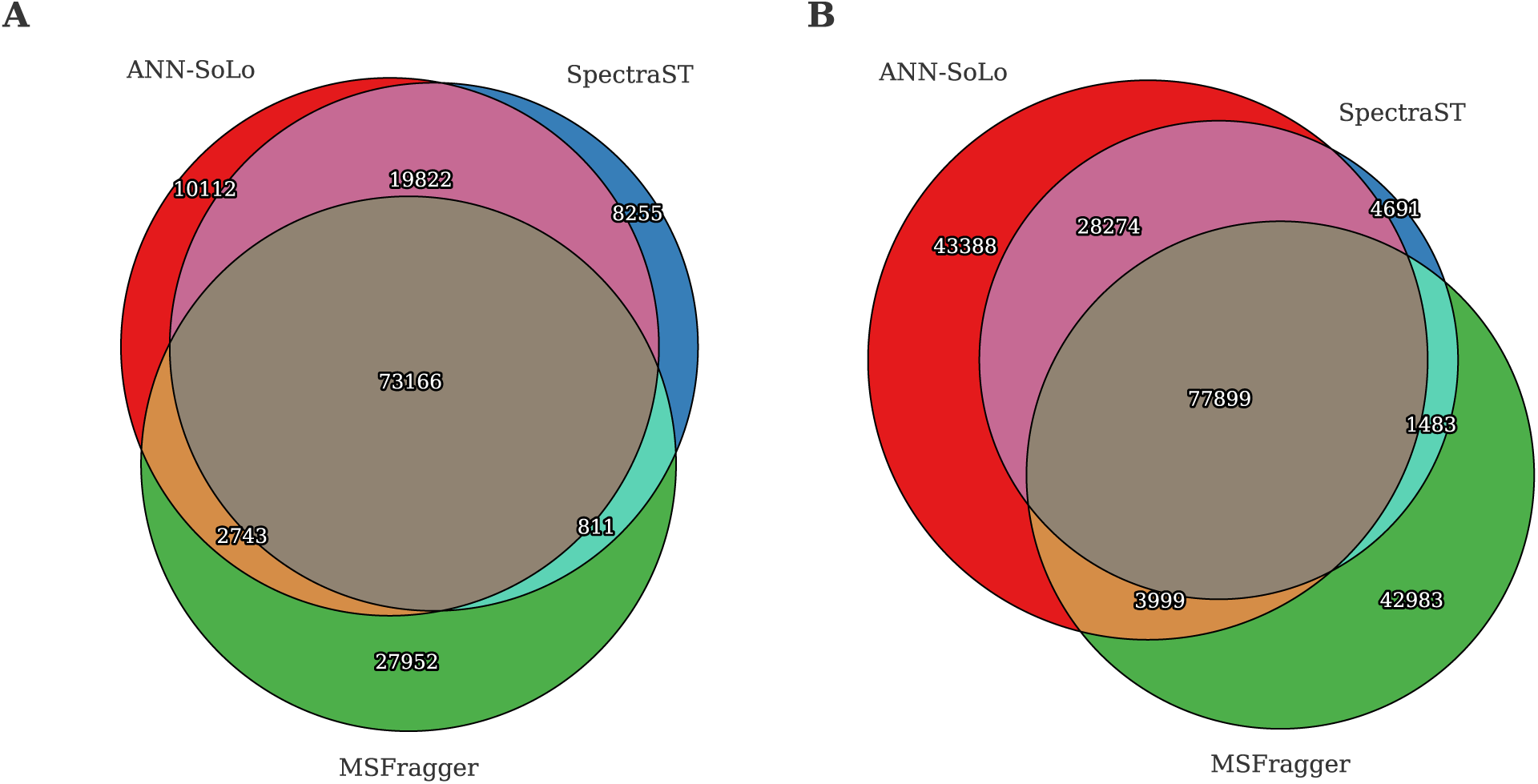
Comparison of identified peptides between ANN-SoLo, SpectraST, and MSFragger on the HEK293 data set. (A) Standard search. (B) Open search.

Next, we have investigated which modifications frequently occur in this HEK293 human cell line data set based on the precursor mass differences for the identified SSMs (supplementary figure S5). By referencing the observed precursor mass differences against the Unimod public database of protein modifications [16] we can derive the chemical events that likely explain the observed mass differences (table 3). We can see that common modifications, such as oxidation, frequently occur. Additionally, mass shifts corresponding to various amino acid substitutions can be frequently observed. These can potentially indicate single amino acid variants, but can likely also be explained by the incomplete coverage of the human proteome by our spectral library. Consequently, OMS can to some extent alleviate a longstanding criticism of spectral library searching in that it is only able to re-identify spectra that have been previously observed. Finally, the precise mapping of these mass differences to a known modification further substantiates the validity of these identifications and confirms that OMS can be used to accurately identify modified peptides.

**Table 3:** The most frequent precursor mass differences for the HEK293 data set and likely modifications sourced from Unimod corresponding to these precursor mass differences. The delta-mass column contains the median precursor mass difference of that SSM subgroup.

| # SSMs | $\Delta m$ (Da) | Potential modification |
| --- | --- | --- |
| 369 341 | 0.003 |  |
| 46 659 | 1.005 | First isotopic peak |
| 31 473 | 15.998 | Oxidation or hydroxylation<br>/ Ala → Ser substitution /<br>Phe → Tyr substitution |
| 14 088 | 2.006 | Second isotopic peak |
| 6 418 | -0.991 | Amidation |
| 5 777 | -17.023 | Pyro-glu from Q / loss of ammonia |
| 5 433 | 17.002 | Replacement of proton with ammonium ion |
| 4 815 | 183.039 | Aminoethylbenzenesulfonylation |
| 4 777 | 27.998 | Formylation / Ser → Asp substitution / Thr → Glu substitution |
| 3 266 | 301.991 | Unidentified modification [15, 40] |

## 4 Conclusions

Here we have introduced the ANN-SoLo spectral library search engine. ANN-SoLo uses ANN indexing to efficiently select the most likely candidate matches from a spectral library based on their spectral similarity with the query spectrum. As candidate retrieval using the ANN index only depends on spectral similarity without taking precursor mass information into account this strategy naturally lends itself to OMS. By using the ANN index the number of candidates that need to be evaluated for each query spectrum can be reduced by orders of magnitude, decreasing the time required to perform an open search accordingly. Furthermore, because the number of potential matches that needs to be evaluated is small this opens up the possibility to use more computationally expensive similarity measures to score SSMs without incurring an overly large performance hit. This is exemplified by our use of the shifted dot product, which allows us to accurately match a modified spectrum to its unmodified counterpart.

Thanks to these advances open spectral library searching has become a feasible strategy for the sensitive identification of modified peptides. We have demonstrated how an extremely large spectral library can be used to detect peptide modifications at a large scale, which can give important insights into their biological activity. Notably, we have used a repository-wide spectral library which has been derived from a massive amount of publicly available spectral data. Using a spectral library of such size for open searching with a traditional search engine would drastically suffer from excessive runtimes. In contrast, ANN-SoLo makes it possible to perform such searches in a reasonable time frame.

The application of ANN indexing need not be restricted to open spectral library searching. Other identification tasks which exhibit a large search space, such as metaproteomics [47, 51], or activities which consist of large-scale spectral processing tasks, such as spectral clustering [24, 31, 32, 61], can similarly benefit from ANN indexing to achieve substantial speedups.

Furthermore, although here we have used an ANN index consisting of an ensemble of random projection trees several alternative methods for ANN indexing exist. Some examples of ANN indexing techniques which have exhibited excellent empirical performance [5] include hierarchical navigable small world graphs [44], LSH [4], product quantization [37], etc. Additionally, non-metric space indexing can potentially be used to retrieve candidates based on the shifted dot product similarity directly [10]. An investigation into whether these techniques are suitable to speed up spectral library searching remains as future work. Another promising approach to achieve further speedups is by making use of specialized hardware such as GPUs, both for candidate selection using an ANN index [36] as well as to evaluate SSMs [6].

Finally, the FDR procedure plays an important role in evaluating the identification results of an open search. As using a very wide precursor mass window leads to a large increase in search space the probability of having a high-scoring spurious match is considerably higher for an open search compared to a standard search, and such high-scoring decoy matches can have a large influence on the number of accepted identifications when using a global FDR strategy. In contrast, the subgroup FDR procedure calculates the FDR separately for spectra that have distinct modifications [27]. In practice we have observed that subgroups that unambiguously correspond to known modifications often contain very few decoy matches. In contrast, many decoy matches do not belong to a common subgroup as their precursor mass difference is randomly distributed across the range of the precursor mass window. Instead these decoy SSMs are combined in the residual group for FDR calculation, minimizing their negative influence on the accepted identifications. Caution has to be observed though as the actual FDR might be underestimated when too small groups are used [27]. It is a standing research question whether alternative approaches are needed for the accurate FDR estimation of open searches [20, 40].

The ANN-SoLo spectral library searching engine is freely available as open source. The source code and detailed instructions can be found at https://github.com/bittremieux/ANN-SoLo.

## Acknowledgement

We want to thank Henry Lam for updating the parsing functionality of NIST spectral libraries in SpectraST and Ruedi Aebersold and Ludovic Gillet for providing the high-resolution yeast spectral library.

W.B. is a postdoctoral researcher of the Research Foundation – Flanders (FWO). W.B. was supported by a postdoctoral fellowship of the Belgian American Educational Foundation (BAEF). W.S.N. was supported by National Institutes of Health award R01 GM121818.

The computational resources and services used in this work were provided by the VSC (Flemish Supercomputer Center), funded by the Research Foundation – Flanders (FWO) and the Flemish Government – department EWI.

## Supporting information

Supplementary figure S1: Sequence similarity between ANN-SoLo and the iPRG2012 consensus results. Supplementary figure S2: Cascade open search versus direct open search iPRG2012 correctness. Supplementary figure S3: Sequence similarity between the iPRG2012 cascade open search and direct open search. Supplementary figure S4: Timing profiling brute-force versus ANN. Supplementary figure S5: HEK293 precursor mass differences. Supplementary table S1: ANN-SoLo hyperparameter evaluation on the iPRG2012 data set.

